# Semi-automated reconstruction of glomerular architecture from 3D confocal microscopy data

**DOI:** 10.64898/2026.07.03.736410

**Authors:** Yoseph M Loyd, Sharon E. Chase, Mira Krendel

**Affiliations:** Dept. of Cell and Developmental Biology, SUNY Upstate Medical University, Syracuse NY 13210

## Abstract

Nephrons are the functional units of the kidney; within each nephron, the glomerulus is the initial site of selective filtration that allows removal of waste products while preserving proteins in the bloodstream. Each glomerulus consists of a network of capillaries surrounded by specialized epithelial cells, podocytes, which mediate selective filtration. Abnormalities in glomerular structure impair renal function, resulting in proteinuria and kidney disease. Although several microscopy-based approaches exist to characterize glomerular architecture and structural abnormalities, quantitative analysis is often limited by labor-intensive image segmentation. In this study we present a semi-automated approach for segmentation and analysis of glomerular architecture from three-dimensional confocal microscopy data. Using mTmG transgenic mice that express membrane-associated EGFP in podocytes and membrane-associated tdTomato across all other cell types, we reconstruct podocyte processes and glomerular capillaries from volumetric renal images. This semi-automated approach reduces manual segmentation effort and supports more efficient, standardized analysis of glomerular architecture in three-dimensional confocal microscopy datasets.

## Introduction

The filtration units of the kidneys, nephrons, are responsible for filtering waste from circulating blood. This selective filtration process takes place in the glomerulus, a complex network of capillaries surrounded by specialized epithelial cells, podocytes (Scott and Quaggin, 2015). Selective filtration by the glomerulus is facilitated by the presence of thin podocyte extensions, called foot processes, which interlace with those of neighboring podocytes to form a sieve-like meshwork. The spaces between adjacent foot processes, known as filtration slits, are bridged by specialized protein assemblies known as the slit diaphragm complexes (Birtasu et al., 2025; Quaggin and Kreidberg, 2008; Scott and Quaggin, 2015). Slit diaphragm complexes are composed of transmembrane proteins, including nephrin, that form cell-cell junctions between foot processes, providing a selective barrier that helps prevent large protein molecules from leaking into the urinary filtrate. Selective glomerular filter components include slit diaphragms and the glomerular basement membrane (GBM) that surrounds glomerular capillaries (Birtasu et al., 2025; Bulow and Boor, 2019; Quaggin and Kreidberg, 2008; Ruotsalainen et al., 1999). The integrity of slit diaphragms and the delicate elongated shape of foot processes are maintained by the podocyte actin cytoskeleton and associated regulatory and scaffolding proteins such as ZO-1, synaptopodin, and podocin in podocytes (Holthofer, 2007; Ichimura et al., 2003; Itoh et al., 2014; Kawachi and Fukusumi, 2020; Mundel et al., 1997). When foot process structure is disrupted, they become effaced (flattened), resulting in disruption of selective glomerular filtration and proteinuria (Kawachi and Fukusumi, 2020; Lee et al., 2020; Pollak et al., 2014). Proteinuria, if not reversed, is associated with more severe changes such as focal and segmental scarring that can be observed by histology; this condition is known as focal segmental glomerulosclerosis (FSGS) (Deegens et al., 2008; Trautmann et al., 2018). Interestingly, foot process effacement is associated with a loss of slit diaphragm and cytoskeletal proteins such as nephrin (Lee et al., 2022) and myosin 1e (Myo1e) (Chase et al., 2012; Krendel et al., 2009).

Electron microscopy (EM) is commonly used to examine healthy and effaced foot processes (Ichimura et al., 2003; Itoh et al., 2014; Krendel et al., 2009; Li et al., 2019; Potter et al., 2019; Randles et al., 2016) and can be used for tissue assessment in pathology (Qasim et al., 2025). Additionally, fluorescence super-resolution optical microscopy of slit diaphragm proteins and foot processes (Siegerist et al., 2018; Suleiman et al., 2017; Unnersjo-Jess et al., 2023) has provided insights into the size and periodicity of foot processes with EM-like resolution. Interestingly, key features of podocyte morphology, including foot processes and their higher-order organization, are readily observed in native kidney tissue but remain challenging to reproduce *in vitro*, as current *in vitro* podocyte culture methods do not fully recapitulate the glomerular microenvironment (Agarwal et al., 2021; Shankland et al., 2007). Although substantial progress has been made in developing advanced substrates, creating matrices that match glomerular mechanical properties, and using micropatterned substrates and microfluidic platforms that partially restore podocyte differentiation and slit diaphragm protein expression (Ashammakhi et al., 2018; Korolj et al., 2018; Poudel et al., 2025; Ron et al., 2017; Tuffin et al., 2019; Wang et al., 2022), there remains a critical need for quantitative, three-dimensional benchmarks of native podocyte architecture against which engineered systems can be evaluated.

In this work, we present a pipeline of image processing steps that enables the semi-automated segmentation of podocyte membrane digitations and glomerular capillaries from three-dimensional confocal datasets. Using confocal imaging of Podo-Cre^+/Tg^ mice crossed with the ROSA26^(ACTB-tdTomato,-EGFP)Luo/J^ reporter mouse strain that exhibits membrane-localized EGFP (mGFP) in cells expressing Cre recombinase and membrane-localized tdTomato (mtdTomato) across all other cell types, we reconstruct the 3D glomerular architecture through a series of image processing steps. This pipeline utilizes an adaptation of the dynamic Bernsen binarization method (Eyupoglu, 2017; Roy et al., 2014) to enable the 3D reconstruction and visualization of foot processes and glomerular capillary segments in fluorescent tissue.

This approach establishes a potentially quantitative structural reference for future comparisons with *in vitro* models and offers a low-barrier method to reduce manual effort in the analysis of glomerular organization. This approach may support the development of 3D *in vitro* culture environments that better reproduce podocyte morphology observed *in vivo* and resemble the images obtained from dual-fluorescent glomerular tissue.

## Results

### Dual fluorescent renal tissue permits the visualization of glomerular structure

Previously, we have established a podocyte-specific Myo1e-KO mouse model to study the role of a component of the podocyte cytoskeleton, myosin 1e (Myo1e), in podocyte functions (Chase et al., 2012). This mouse model uses Cre recombinase expression under the control of the podocin promoter (Moeller et al., 2003) to selectively perform Cre-mediated recombination in podocytes. The Cre-lox system can also be used to selectively express mGFP in podocytes. In this work we crossed the Podo-Cre^+/Tg^ mice with the ROSA26^(ACTB-tdTomato,-EGFP)Luo/J^ mice, also known as mTmG mice, which express membrane-targeted red fluorescent protein, mtdTomato, prior to Cre-mediated recombination and membrane targeted EGFP, mGFP, after recombination (Muzumdar et al., 2007). This allowed us to generate dual fluorescent mice exhibiting mGFP expression in podocytes and mtdTomato across all other cell types. Since the junctional protein ZO-1 is more highly expressed in podocytes/glomeruli than in the tubulointerstitium, we used immunostaining with anti-ZO-1 antibodies on cryosections of renal tissue from dual fluorescent mice, to verify that most renal cells (such as tubular epithelium, endothelium, and stromal cells), subsequently referred to as “tissue”, express mtdTomato, and specialized glomerular cells, podocytes, exhibit mGFP expression (Figure 1).

**Figure 1.**
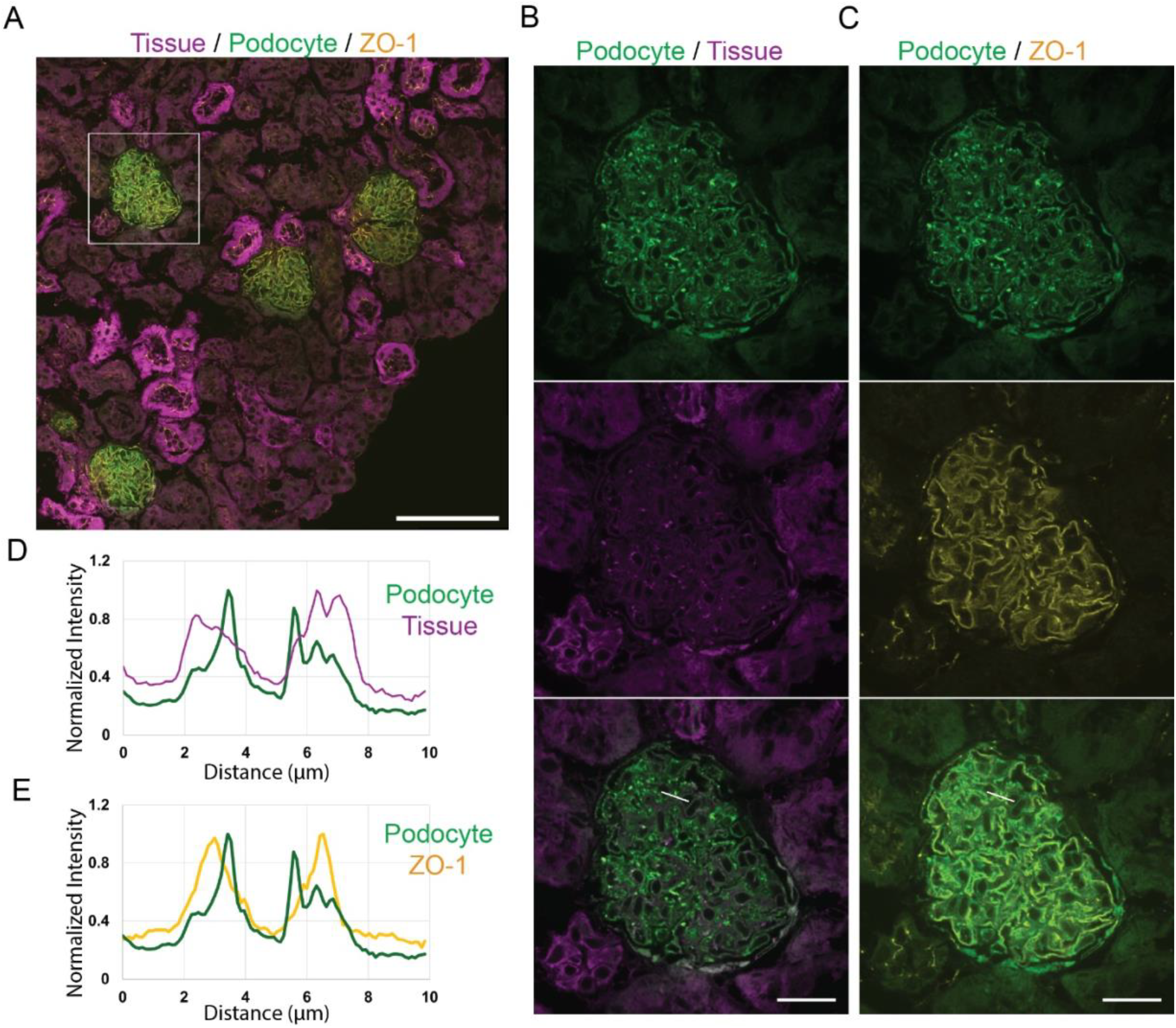
Podocyte-specific expression of mGFP permits glomerular imaging. (A) Confocal section of a paraformaldehyde-fixed cryosection of kidney cortex from podocyte-mTmG reporter mouse stained with antibodies against ZO-1. mtdTomato fluorescence marks all renal structures (tubular, vascular, and stromal, labeled as “tissue”), while mGFP labels podocyte membranes. Scale bar, 100 µm. (B) Region of interest (ROI) indicated by the white outline in panel A shows a glomerulus and surrounding tubules. Scale bar, 20 µm. (C) Same ROI illustrating GFP-labeled podocyte membranes and ZO-1 immunofluorescence. Scale bar, 20 µm. (D, E) Line-scan analysis of panels B and C, respectively. (D) Graphs display normalized fluorescence intensity of the mGFP (podocytes) and mtdTomato (tissue) signals along the white line shown in the merged image in Panel B. (E) Graphs display normalized fluorescence intensity of podocyte membranes and ZO-1 signal along the white line shown in the merged image in panel C.

High-resolution imaging of individual glomeruli revealed that mGFP-expressing podocytes closely neighbor mtdTomato-expressing cells that likely comprise glomerular capillary networks or mesangial cell clusters (Figure 1B, D). Immunostaining for the tight junction and slit diaphragm-associated protein ZO-1 showed strong enrichment in the glomerular regions overlapping with the areas of podocyte mGFP signal (Figure 1C, E). This confirms the selective expression of mGFP in podocytes.

We next examined whether thick unfixed sections of renal tissue from podocyte-mGFP mice enable three-dimensional confocal imaging of glomeruli for visualization of podocyte morphology. Volumetric confocal datasets visualized in UCSF ChimeraX (Goddard et al., 2018; Pettersen et al., 2021), and ImageJ/FIJI (Schindelin et al., 2012) revealed mGFP-labeled podocyte membranes in both volumetric maximum intensity projections (vMIPs) and orthogonal planar views (Figure 2). These 3D reconstructions revealed striated patterns within the glomerulus, likely corresponding to the podocyte major processes and foot processes.

**Figure 2.**
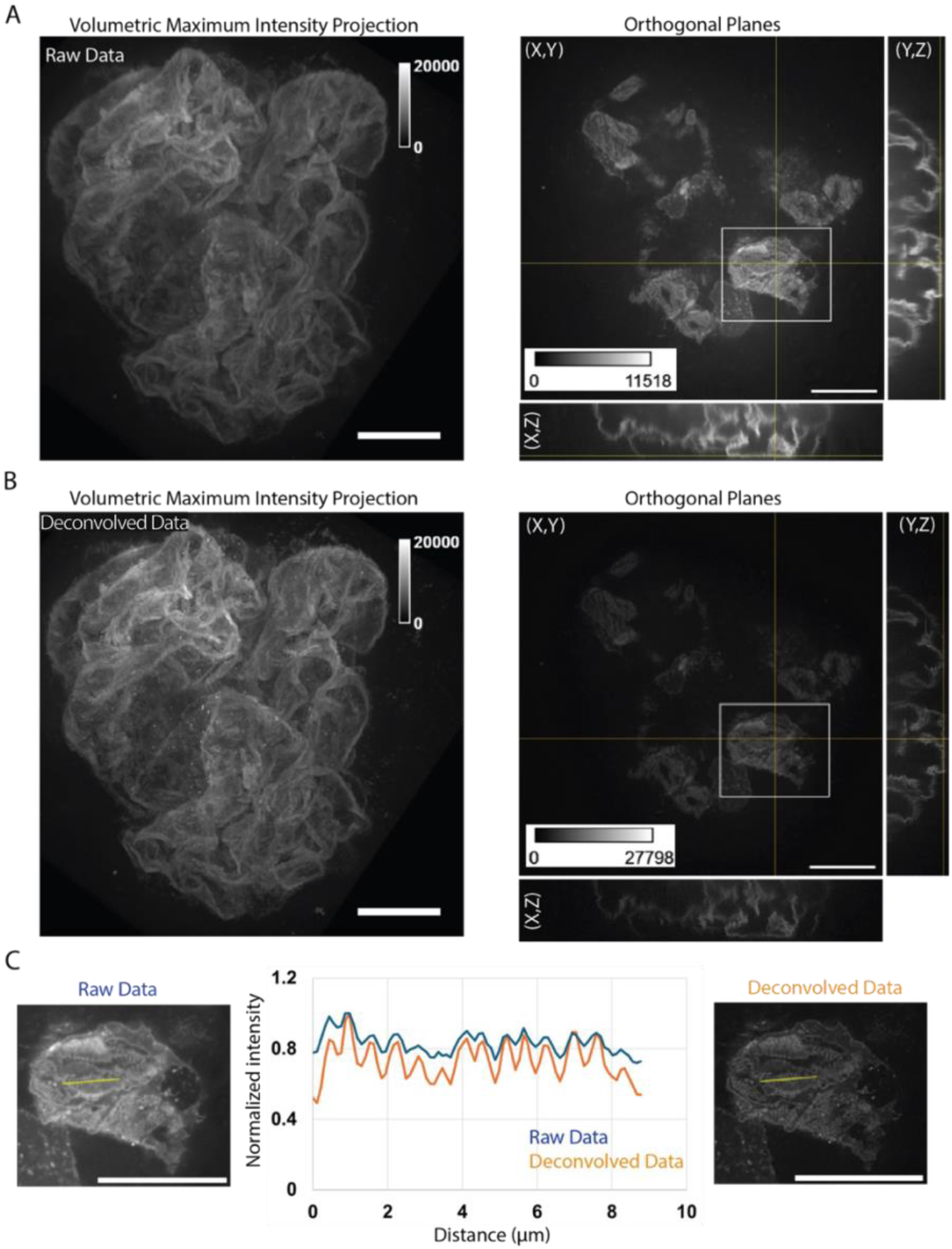
Visualization of digitated podocyte processes in 3D images of renal tissue through targeted mGFP expression and image processing. (A) Raw 3D confocal imaging data of podocyte membranes from manually sectioned renal tissue from a podocyte-mTmG reporter mouse. Data are displayed as either a vMIP or orthogonal planar views. Grayscale intensity bars are displayed for each image set. Scale bar, 20 µm. (B) The result of deconvolution of the 3D confocal imaging dataset of podocyte membranes from panel A. Data are visualized as both a vMIP and orthogonal planar views. Grayscale intensity bars are displayed for each image set. Scale bar, 20 µm. (C) Single confocal sections (raw and deconvolved) of the regions indicated by the square white ROIs in panels A and B. Line-scan analysis of raw and deconvolved images showing normalized intensity profiles of podocyte membranes along the yellow lines shown in each confocal section. Scale bars, 20 µm.

Although the initial images exhibited a low signal-to-noise ratio, striated patterns of podocyte processes became apparent after deconvolution of the 3D confocal datasets to remove out-of-focus signal (compare Figure 2A and 2B). Line-scan analysis of mGFP intensity along the yellow lines within the white ROIs (Figure 2C) highlights peaks of fluorescence signal at approximately half-micron intervals in both raw and deconvolved data, with the more distinct peaks and valleys in the deconvolved images. These regularly spaced intensity peaks are consistent with membrane digitation, likely reflecting the organization of podocyte major and foot processes along glomerular capillaries.

### Development of a 3D semi-automated pipeline to segment glomerular structure

Having shown that confocal microscopy imaging is sufficient to resolve podocyte membrane digitation in images of fresh unfixed tissue, we proceeded to develop a pipeline for the automatic segmentation of podocyte processes in fluorescence data. As an example, using three regions of interest (ROIs) of a deconvolved image of mGFP-labeled podocyte membranes in unfixed tissue (Figure 3A) we perform the steps outlined in Figure 3B to identify podocyte membrane digitations.

**Figure 3.**
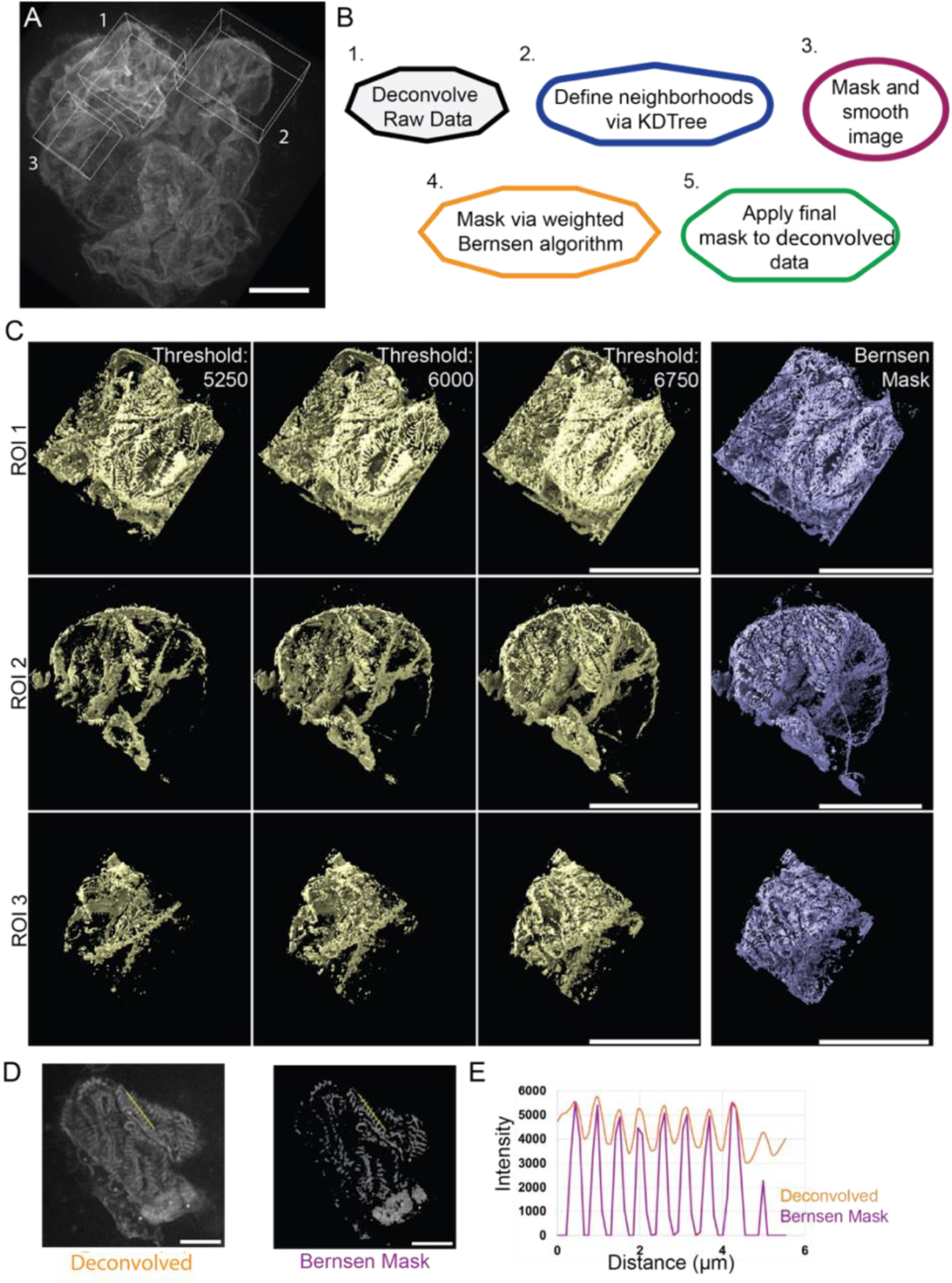
Implementation of a modified 3D Bernsen local thresholding algorithm for segmentation of fluorescent podocyte processes. (A) vMIP of a 3D confocal dataset of mGFP-labeled podocyte membranes from manually sectioned renal tissue from a podocyte-mTmG reporter mouse, with 3 ROIs indicated by white boxes. Scale bar, 20 µm. (B) A simplified stepwise flowchart for the automated segmentation workflow for identification of processes from the confocal datasets. (C) ROIs indicated in panel A are processed using global intensity thresholds (with threshold values indicated) or weighted Bernsen thresholding (a dynamic local thresholding method). Scale bars, 20 µm. (D) A single z plane of digitated podocyte membranes is displayed either as the deconvolved data or the data processed via the Bernsen method. Scale bar, 10 µm. (E) Line-scan analysis along the yellow lines shown in panel D, comparing the intensity profiles of the deconvolved data and Bernsen-processed data.

First, the data are deconvolved before any subsequent processing, ensuring a higher signal-to-noise ratio or higher quality image. Next, to analyze local patterns, a spatial indexing method (KD tree (Bentley, 1975)) is used to identify each pixel’s neighbors within a 3-pixel radius, subsequently referred to as pixel neighborhoods. Defining pixel neighborhoods is essential for the later application of dynamic or local thresholding (the weighted Bernsen method) (Eyupoglu, 2017; Roy et al., 2014). The input image then undergoes background subtraction where I_bg_ is the background-subtracted image, I_i_ is the initial deconvolved image and σ is the standard deviation of intensity values in the deconvolved image, equation 1

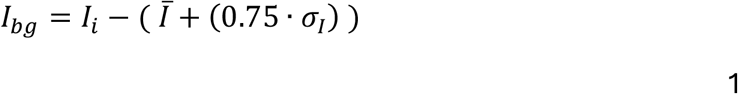

The output image, I_bg_, is then smoothed and masked for all values above background. Once this preprocessing is done, we then implement a 3D Bernsen algorithm for dynamic thresholding. Dynamic thresholding adjusts the signal-to-background cutoff for each pixel based on the local intensity profile of its surrounding neighborhood, improving regional masking compared with a single global threshold. The implementation of the 3D Bernsen algorithm with weights for dynamic thresholding is performed where global contrast is defined as Contrast_g_ that is equal to the mean intensity of all pixels in the image, equation 2

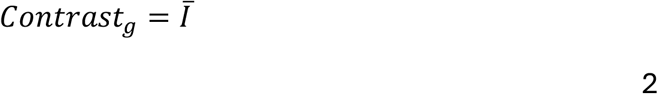

Neighborhood contrast is defined as Contrast_l_ that is a function of I_max_ , the maximum intensity of pixel values in the pixel neighborhood, and I_min_ , the minimum intensity of pixel values in the pixel neighborhood, equation 3

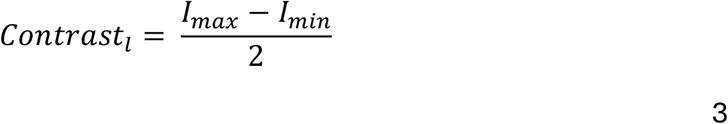

And neighborhood threshold, Thresh, is defined as a function of the maximum intensity of pixel values in the pixel neighborhood, I_max_, the standard deviation of pixel intensities in the pixel neighborhood, σ_IL_, and user-defined constants α and β, equation 4

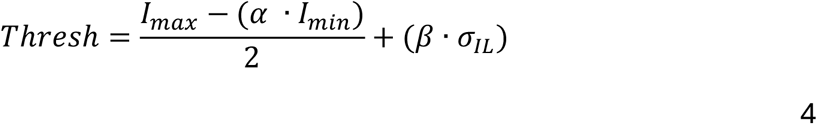

Once global contrast, neighborhood contrast, neighborhood threshold were calculated, we applied the following weighted Bernsen algorithm, where Bernsen Mask_X,Y,Z_ is the resulting binarized image generated using the weighted Bernsen algorithm, Contrast_l_ is the neighborhood contrast, Contrast_g_ is the global contrast, I_X,Y,Z_ is the intensity of a pixel corresponding to position (X,Y,Z), Thresh is the neighborhood threshold, ρ and τ are user defined constants, equation 5

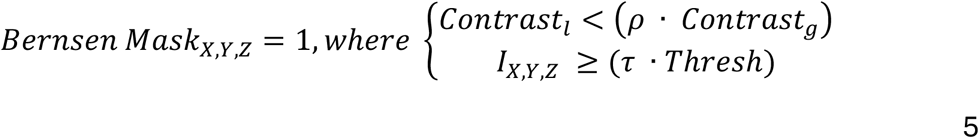

Comparison of the binary mask generated using this dynamic method to simple global thresholding shows that the dynamic thresholding approach better captures podocyte membrane digitations (Figure 3C). While simple thresholding could be performed manually on specific subregions to achieve a similar segmentation output, doing so on every subregion of the data is laborious and non-standardized. Using the modification of the Bernsen thresholding algorithm simplifies the thresholding process and allows automated generation of a mask for 3D rendering of podocyte processes (Figure 3).

We have also compared the weighted Bernsen method for podocyte segmentation with creating a binary mask using the Trainable WEKA Segmentation tool, a machine learning method that uses manual user input to train the segmentation plugin (Arganda-Carreras et al., 2017). While both segmentation approaches identify podocyte membranes, the WEKA segmentation tool fails to generate a 3D surface reconstruction resembling the deconvolved data or podocyte digitated membranes (Figure 4A, B). Line-scan analysis in two different ROIs with visible podocyte digitations shows that the WEKA segmentation tool did not improve segmentation of bright ROIs from the initial deconvolved data (Figure 4 C, D). On the other hand, line-scan analysis revealed that the Bernsen method retained intensity peaks corresponding to the peaks initially identified from the deconvolved images while excluding noisy spaces between digitations (Figure 4C, D).

**Figure 4.**
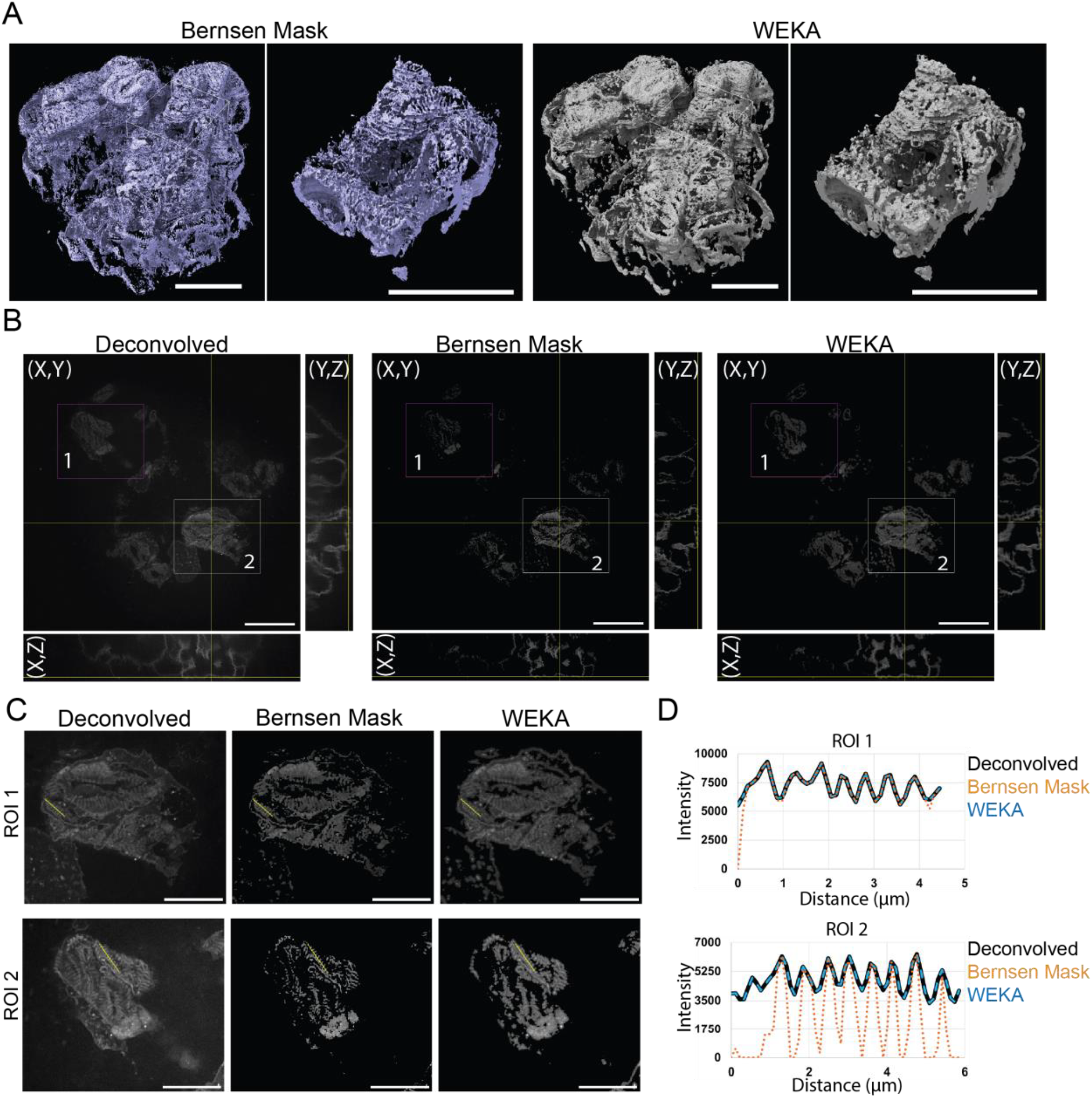
3D implementation of the weighted Bernsen algorithm permits segmentation of podocyte processes in tissue with podocyte-targeted membrane fluorescence. (A) The 3D surface representation of the weighted Bernsen algorithm’s mask for local thresholding and WEKA segmentation mask of podocyte membranes. Scale bars, 20 µm. (B) Confocal optical sections of podocyte membranes in a nephron displayed as orthogonal planes (XY, XZ, and YZ). Shown are the planar views of the deconvolved dataset, the deconvolved dataset masked using the weighted Bernsen method, and the deconvolved dataset masked using the WEKA segmentation approach. Scale bar, 20 µm. (C) ROIs shown in panel B are enlarged to display the deconvolved data and the masks obtained using the weighted Bernsen algorithm and the WEKA segmentation tool. Scale bar, 10 µm. (D) Line-scan analysis along the lines shown in panel C, comparing the deconvolved data and outputs of the masks obtained using our implementation of the weighted Bernsen algorithm and the WEKA segmentation tool.

### Dynamic thresholding provides the opportunity to interrogate glomerular capillary networks

To enable automated quantification of glomerular capillary morphology, we combined the modified weighted Bernsen segmentation pipeline with glomerular isolation by sieving. Dual fluorescent renal tissue was sieved as described in Methods to isolate intact glomeruli, allowing imaging of physically separated nephron structures where capillaries remain surrounded by podocytes (Figure 5A). Capillary reconstruction follows eq 1-4 from the previously described weighted Bernsen thresholding with a minor alteration to eq 5. By inverting the previous contrast gating while selecting mtdTomato channel, we can mask capillary networks, equation 6.

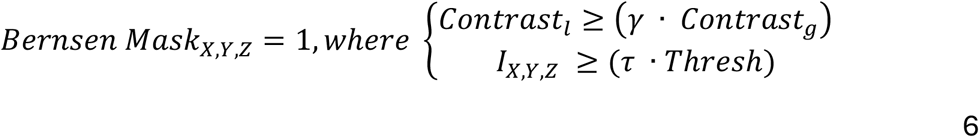

**Figure 5.**
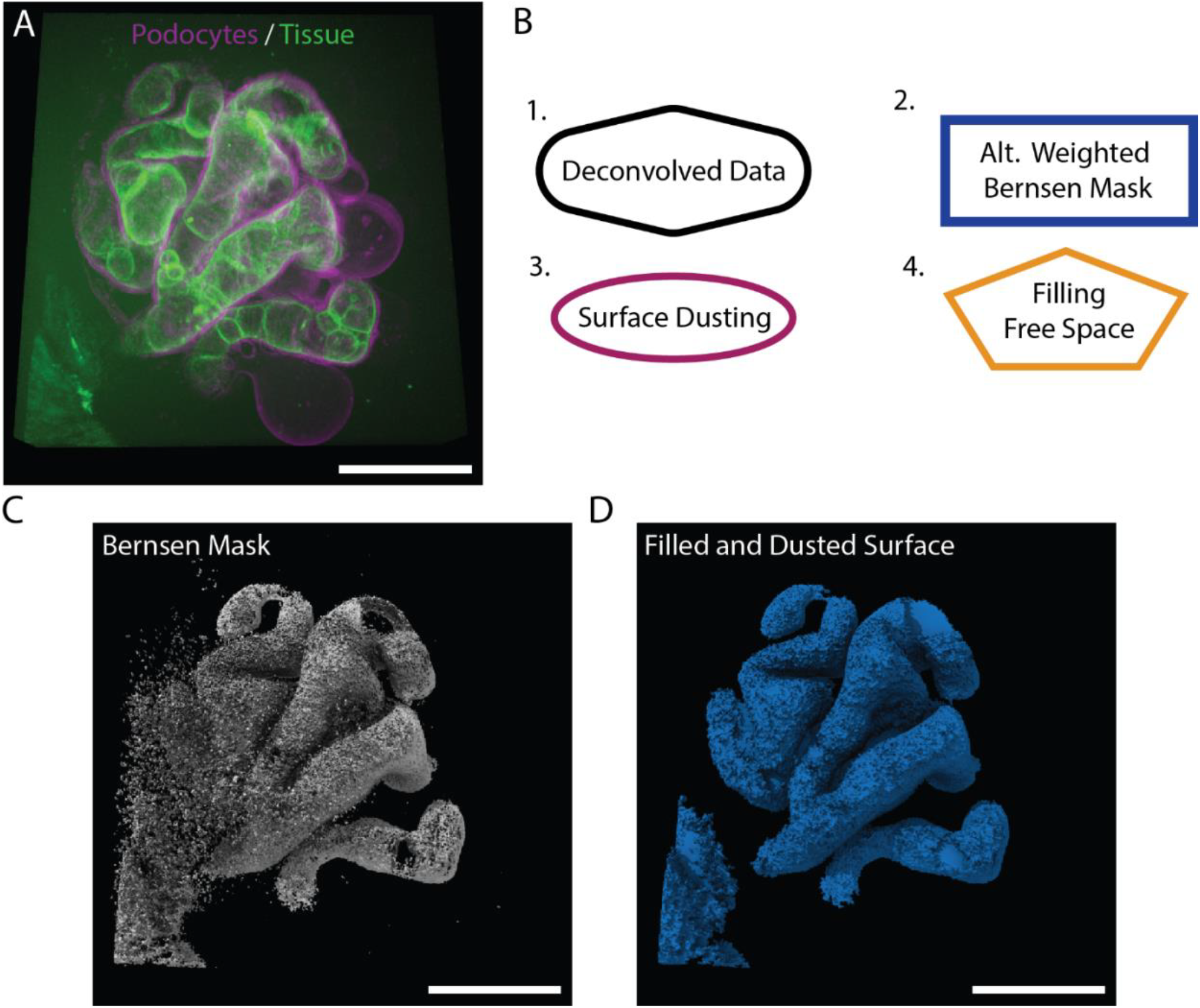
Reconstruction of glomerular capillaries from mtdTomato expressing renal tissue. (A) A vMIP representation of the 3D confocal dataset of fresh unfixed tissue isolated by sieving from the podocyte-mTmG reporter mouse. Membrane fluorescence of endothelial cells expressing mtdTomato permits the visualization of glomerular capillaries while podocytes are visualized via targeted expression of mGFP. (B) Workflow for capillary reconstruction and the initial morphological analysis in subsequent panels. (C, D) 3D surface representations generated from the workflow outlined in panel B. Included are the initial mask of capillaries from the weighted Bernsen thresholding (C) and the surface-dusted and hole-filled capillary mask (D). Scale bars, 20 µm.

The initial glomerular capillary masks produced by this pipeline contained substantial noise, and many sections of reconstructed capillaries had holes and discontinuities in the reconstructed surface (Figure 5C). To correct for these imperfections in the reconstructed surface, we applied a series of post-processing operations, including a surface dusting operation to remove small, isolated noise elements, an area-closing step (Vincent, 1993) to fill small holes in the surface, and a free space identification step to attempt to fill large holes (Figure 5B,D). Although these steps improved surface continuity, residual imperfections remained and were manually corrected when necessary (e.g. clipping the glomerular basal plane to fully disconnect neighboring capillaries).

## Discussion

In this work, we report the development of a workflow that allows users to visualize and segment three-dimensional structures in dual-fluorescent tissue using confocal microscopy and semi-automated image analysis based on dynamic local image thresholding. This method is anticipated to enable observation and quantification of morphological features in complex tissue microenvironments and organoid models. For the purposes of the current study, we focused on the initial conceptual development of the image analysis pipeline through semi-automated segmentation of glomerular structures in dual-fluorescent reporter mice that express mGFP in podocytes and mtdTomato in other cell types. Having validated podocyte-specific mGFP expression by co-immunostaining of cryosections of renal tissue for ZO-1, which is enriched in podocytes, we subsequently acquired three-dimensional confocal images of glomeruli from mouse kidney tissue sectioned or sieved without fixation. Resulting images of podocyte membranes displayed approximately half-micron-wide digitated structures in subregions of the glomerulus. While all podocytes are expected to express mGFP, differences in the amount of mGFP expressed in individual podocytes likely allow observation of these striated patterns. More reliable identification of podocyte processes throughout the kidney may be achieved in the future by using inducible expression of Cre recombinase with low level of induction treatment, for example a brief pulse of doxycycline, to increase the heterogeneity of mGFP expression. The width of the glomerular striations observed using confocal microscopy resembles the spacing and size of foot processes that have been observed in mouse tissue using super-resolution fluorescence microscopy and electron microscopy, with the podocyte foot process width ranging from 250 nm to 500 nm in healthy adult mice (Basgen et al., 2021; Siegerist et al., 2017; Unnersjo-Jess et al., 2023; Veron et al., 2010; Zhong et al., 2018). Similarly, in our previous papers we have observed podocyte foot process density by TEM to be approximately 1.5 – 1.6 foot processes per micrometer, which corresponds to a spacing of approximately 0.6 – 0.7 µm, broadly consistent with the width of periodic membrane patterns observed in this study (Chase et al., 2012; Krendel et al., 2009).

Deconvolution of confocal datasets to remove systematic blurring inherent to optical microscopy (Sibarita, 2005) resulted in improved image quality. However, both raw and deconvolved images exhibited uneven brightness, which increased visual complexity and made it more difficult to present observed structure in volumetric maximum-intensity projections. Attempts to reduce this visual complexity and isolate podocyte foot processes by performing simple (global) thresholding failed to resolve podocyte morphology across the glomerulus due to uneven brightness. Thus, as an alternative to simple thresholding, we implemented an adaptation of a three-dimensional dynamic Bernsen binarization/thresholding algorithm for semi-automated segmentation of podocyte processes and glomerular capillary topology. This modified weighted Bernsen method presented in this paper showed greater sensitivity for identifying bright local ROIs compared with our attempts to use the WEKA segmentation tool. The corresponding isosurface renderings of the Bernsen-derived masks revealed three-dimensional architecture of podocyte processes by confocal microscopy that resembled three-dimensional reconstructions of podocytes obtained by FIB-SEM (Ichimura et al., 2015). Together, these results demonstrate that a rapid segmentation approach can be used to observe podocyte three-dimensional morphology without the need for super-resolution microscopy or electron microscopy.

Building on our ability to segment membrane digitations corresponding to podocyte processes, we introduced a modification to the pipeline that enabled reconstruction of glomerular capillaries based on mtdTomato expression. Although still preliminary, this approach generates masked capillary networks that can serve as a foundation for future analyses of three-dimensional glomerular architecture. In particular, these masks may be useful for defining capillary connectivity and for measuring both global and local capillary diameters.

We anticipate that in the future this segmentation approach would be more successful if utilized for podocyte segmentation in images collected from Confetti mice expressing multi-color podocyte reporters to observe interdigitation of foot processes in glomeruli and morphologic characteristics of individual podocytes (Tao et al., 2014). Doing so would additionally permit quantification of foot process effacement and irregularities in the spatial patterns of podocyte processes in mouse models of disease without the need for super-resolution or electron microscopy techniques. Additionally, while our pipeline performs well overall, object segmentation can be further improved by increasing the number of processing layers involved, which would help transition this semi-automated approach to a fully automated method. In future iterations we envision that a fully automated pipeline would permit the complete processing of several collections of three-dimensional glomerular datasets per day to reduce user labor and increase processing throughput. Once adequate segmentation is achieved, quantitative output can be further refined to describe local morphologic characteristics of glomerular capillaries and podocyte processes. In addition to advancing studies of kidney disease progression *in vivo*, morphological information generated by this pipeline may also inform the design and fabrication of artificial glomerular-like substrates for *in vitro* podocyte culture. Overall, we present a broadly applicable segmentation framework that enables reconstruction of podocyte foot processes and glomerular capillaries from dual-fluorescent renal tissue, providing an accessible platform for quantitative analysis of glomerular architecture.

## Methods

### Animals and genotyping

To obtain podocyte-mGFP/mtdTomato (Podo-Cre mTmG) reporter mice, we used mice from our colony that were a cross between Myo1e^Flox/Flox^ or Myo1e^WT^ or Myo1e^-/-^ mice and Podo-Cre Rosa^mTmG^ transgenic mice. Myo1e^-/-^ and Myo1e^Flox/Flox^ mice have been previously described (Chase et al., 2012) and were maintained on the C57/BL6 background. NPHS2-Cre (Podo-Cre) mice (Moeller et al., 2003) were also maintained on the C57/Bl6 background in our lab. mTmG mice, B6.129 (Cg)-Gt (ROSA)26Sortm4 (ACTB-tdTomato,-EGFP) Luo/J, (stock# 007676) were purchased from Jackson Labs. Most mice used in this paper had at least one WT allele of Myo1e and exhibited no detectable albuminuria as verified using SDS-PAGE of urine samples; thus, these mice were used as phenotypically normal reporter mice. All mice used were heterozygous for Podo-Cre and mTmG transgenes. All protocols have been approved by SUNY Upstate Medical University’s IACUC under protocol# 364.

Genotyping for Myo1e and Podo-Cre was performed as previously described (Chase et al., 2012). Primers mTmG#1 (CTCTGCTGCCTCCTGGCTTCT), mTmG#2 (CGAGGCGGATCACAAGCAATA), mTmG#3 (TCAATGGGCGGGGGTCGTT) were used to genotype mTmG mice (the presence of a 330 bp band indicates that mTmG allele is not present, the presence of a 250 bp band designates the presence of the mTmG transgene).

### Kidney preparation

Mice were euthanized in accordance with IACUC-approved procedures. Post euthanasia kidneys were harvested, kidney weight was recorded, and then kidneys were manually bisected. 2 kidney halves were cryopreserved in Tissue-Tek O.C.T. Compound (Sakura FineTek; Torrance, CA) and 1 half was embedded in paraffin. The remaining kidney half was used for immediate imaging, where it was further sectioned into several mm thick tissue samples and immersed in 1x PBS (phosphate buffered saline) in a glass vial at room temperature temporarily. Samples were then loaded into a 35 mm glass bottom dish (Mattek; Ashland, MA) with 500 µL of 1x PBS, gently using forceps to position them for confocal imaging.

### Glomerular isolation by sieving

Harvested kidneys were submerged in 4°C 1x PBS, then stored on ice. A series of sieves was then placed over the top of a beaker in ascending filter sizes, 71 µm, 100 µm, and 180 µm. Harvested kidneys were then loaded onto the 180 µm filter and gently ground through the sieve using the base of a glass beaker as a pestle. The sieve was then gently rinsed with 3 L of 4°C 1x PBS. The 180 µm sieve was then removed, and waste was discarded. The grinding and rinsing process was repeated with the 100 µm sieve. After rinsing the 100 µm sieve, waste was discarded, and the glomerular material was collected on the 71 µm sieve. Tissue was collected from the 71 µm sieve, using 4°C 1x PBS, a wash bottle, and 1ml pipette. The collected tissue was moved to a conical tube and suspended in 4°C 1x PBS. The tissue suspension was centrifuged at 4°C at 1000g for 5 minutes. Supernatant was removed from the conical tube, and tissue was resuspended in 4°C 1x PBS.

### Cryosectioning and immunostaining

O.C.T.-embedded kidney tissue was sectioned at 10 µm thickness using CM1950 cryostat (Leica; Wetzlar, Germany) onto Superfrost Plus slides (Fisher Scientific; Carlsbad, CA). Cryosections were stored at -80°C overnight. The following day slides were left to dry at room temperature for 15 minutes in the dark. Once dried, sections were then rehydrated in 1x PBS for 10 minutes and fixed by adding 500 µl of 4% paraformaldehyde (Electron Microscopy Sciences; Hatfield, PA) in 1x PBS solution and incubating for 15 minutes at room temperature. Slides were then washed 3 times in a Coplin jar filled with fresh 1x PBS for 15 min in the dark at room temperature, and sections were permeabilized by adding 500 µl of a 0.25% Triton-X100 (JT Baker; Radnor, PA, cat# JTB-X198-09) in 1x PBS solution to each sample followed by incubation for 5 minutes at room temperature isolated from light in a humidified box. Once permeabilized, samples were washed 3 times. Non-specific binding was blocked using 500 µl of a 3% (w/v) bovine serum albumin (BSA) (Research Products International; Mount Prospect, IL) solution in 1x PBS at 37°C for 30 minutes in a humidified box in the dark. After blocking, samples were washed 3 times. Primary antibody labeling was performed by adding 250 µl of a 500 µl 1x PBS containing 1 µl of rabbit anti-ZO1 (Thermo Fisher Scientific; Carlsbad, CA, cat#61-7300) and incubating for 1 hour isolated from light at 37°C in a humidified box. After primary labeling, samples were washed 3 times. A secondary antibody labeling was subsequently done by adding 500 µl of 1000 µl 1x PBS containing 1 µl of goat anti-rabbit IgG - AlexaFluor-647 (Jackson Immuno Research; West Grove, PA cat# 111-605-003) and incubating for 1 hr isolated from light at 37°C in a humidified box. After this step samples were washed 3 times, and DNA was stained by adding 500 µl NucBlue fixed cell stain (Thermo Fisher Scientific; Waltham, MA, cat# R37606) solution comprised of 1 drop NucBlue reagent in PBS and then incubating for 30 min isolated from light at 37°C in a humidified box. After incubation, samples were washed 3 times and mounted using Prolong Gold Antifade Mountant (Thermo Fisher Scientific; Carlsbad, CA). Mounted samples were then cured for 24 hr isolated from light at room temperature and stored at 4 C until imaged.

### Data acquisition

Confocal microscopy data was collected on a Ti2E spinning disk confocal microscope (Nikon; Tokyo, Japan), equipped with the CSU-X1 spinning disk (Yokogawa; Tokyo, Japan), a LUNF-XL (Nikon; Tokyo, Japan) laser combiner and an iChrome MLE-LFA-NI2 (TOPTICA Photonics; Munich, Germany) 4 laser line source with 405 nm, 488 nm, 561nm, and 640 nm lasers. Emitted light was passed through the CSU-X1 filter wheel alternating through the following filters for associated laser lines, 405 nm ET455/50M (Chroma; Bellows Falls, VT), 488nm ET525/36M (Chroma; Bellows Falls, VT), 561 nm ET605/52M (Chroma; Bellows Falls, VT), and 640nm ET705/72M EM (Chroma; Bellows Falls, VT). Imaging settings were optimized for minimal photobleaching per kidney section. Sample scanning was fully motorized for X, Y, Z scanning on this instrument and controlled through NIS elements version 5.42.02 (Nikon; Tokyo, Japan). Images were collected using the Nikon CFI APO 60X OIL 1.4NA 0.14MM WD LAMBDA S objective (Nikon; Tokyo, Japan), and an Orca FusionBT camera (Hamamatsu; Hamamatsu, Japan) for image acquisition.

Images were post-processed as follows. Deconvolution of images was achieved through using NIS Elements Advanced Research version 5.42.02 (Nikon; Tokyo, Japan) and subsequently images were visualized using UCSF ChimeraX (Goddard et al., 2018; Pettersen et al., 2021) and ImageJ (Schindelin et al., 2012).

## Acknowledgements

This work was supported by the National Institute of Diabetes, Digestive, and Kidney Diseases awards R01DK083345 and R21DK136083 to MK. Raw data are available on https://figshare.com/projects/Semi-automated_reconstruction_of_glomerular_architecture_from_3D_confocal_microscopy_data/268214

## Author contributions

Conceptualization: Y.M.L. and M.K.; Methodology: Y.M.L., S.E.C., and M.K.; Software: Y.M.L.; Investigation: Y.M.L. and S.E.C.; Animal breeding, genotyping, tissue extraction, and glomerular isolation by sieving: S.E.C.; Formal analysis: Y.M.L.; Visualization: Y.M.L.; Writing - Original Draft: Y.M.L.; Writing - Review & Editing: Y.M.L., S.E.C., and M.K.; Supervision: M.K.; Funding acquisition: M.K.

